# A systematic review of the factors affecting sperm binding and release from the oviduct using *in vitro* experimental animal models

**DOI:** 10.1101/2025.02.03.636345

**Authors:** Muloongo C. Sitali, Madalitso Chelenga

## Abstract

After insemination, spermatozoa bind to epithelial cells of the isthmic part of the oviduct to form a functional sperm reservoir responsible for regulating sperm viability and capacitation before the ovulation signal that triggers its release. Understanding this information is essential to optimizing *in vitro* fertilization (IVF) protocols and contributing to the efforts to develop treatment for infertility resulting from fertilization failure. This review aims to provide a comprehensive and systematic assessment of the literature on the non-steroid hormone factors involved in sperm binding and release from the sperm reservoir in animal species using *in vitro* experiments. PubMed, Web of Science, and Google Scholar databases were searched, and eligible studies included articles published between 1^st^ January 1990 and 31^st^ July 2024 reporting factors involved in sperm binding and release from the oviduct, and this could be used to optimize IVF systems and treatment of fertilization failure-related infertility. Studies that co-cultured sperm and oviductal explants or oviductal epithelial cells from bovine, porcine, canine, camelid, equine, hamster, and ovine were used, and different culture systems were evaluated in this study. Overall, sperm binding to and from the oviductal epithelial cells was influenced by several factors, such as carbohydrates and their derivatives, seminal plasma proteins, glycoproteins, tyrosine phosphorylation of sperm, endocannabinoids, sex-sorting, and glycosaminoglycans. These factors either promoted or inhibited the binding sperm binding to and from oviductal cells. We therefore postulate that these non-steroid factors are potential candidates that can be used to optimize IVF outcomes in both human and animal species.

## Introduction

During copulation, significantly large numbers of sperm are ejaculated into the female genital tract. However, only a few reach the site of fertilization, located in the ampulla region of the oviduct [1]. In most domestic animal species, such as the cow and gilt, a proportion of sperm is stored and can survive in the female reproductive tract for at least 24 to 30 hours after insemination [2]. This feature is attributed to the ability of sperm to form intimate interactions with the oviductal epithelial cells (OECs) to create a reservoir that is located mainly in the isthmus region of the oviduct [2]. Sperm-oviduct interactions help to reduce the chances of polyspermy, maintain sperm’s oocyte-fertilizing competency, and regulate capacitation and hyperactivation [3]. Evidence shows that sperm that fail to bind to the oviductal epithelium, probably due to damage or altered morphology, eventually become immotile and are unable to be released from the reservoir to fertilize the oocyte [4]. This feature suggests that sperm-oviductal interaction is also important for prolonging sperm survival and sperm selection, ensuring that only competent spermatozoa have a chance to fertilize the oocyte [4,5]

It has been reported that, due to the oviduct’s narrow anatomy and its thicker fluid content, sperm movement is slowed down, thereby increasing their contact and binding with the oviductal mucosal epithelium to form a sperm reservoir [4]. It is known that the oviductal environment plays a very critical role in mediating sperm binding and release from the oviductal cells. This includes the action of steroid hormones, notably estrogen and progesterone, whose effect on oviductal function coincides with the estrus cycle [6]. While the effect of steroid hormones on the oviductal function has been comprehensively reviewed elsewhere [7], the same has not been done for non-steroidal counterparts. Evidence shows that sperm-oviductal binding could involve several factors beyond the action of steroid hormones, including the need for carbohydrate recognition for sperm to bind to oviductal cells. Carbohydrate recognition involves the binding of carbohydrates, such as fucose, found on the surface of the oviductal epithelium, to the fucose-binding molecules on the surface of sperm [8]. To date, interest in the research to further understand other factors and events surrounding sperm binding and release from the sperm reservoir has increased. It is envisaged that a comprehensive understanding of these factors would contribute to optimizing the different assisted reproductive technologies (ARTs), especially *in vitro* fertilization (IVF) technology using sexed or non-sexed semen as IVF media mostly excludes non-steroidal hormone factors discussed in the present paper.

Therefore, our systematic review aims to: 1) provide a comprehensive systematic assessment and consolidation of the findings from literature to identify the non-steroid hormone factors involved in sperm binding to and release from the sperm reservoir in animal species, and 2) to provide insights on the possibility to improving IVF and fertility treatment by exploring the application of non-steroidal hormone factors as potential candidates.

## Methods

### Eligibility Criteria

#### Inclusion Criteria

The Preferred Reporting Items for Systematic Reviews and Meta-Analyses (PRISMA) guidelines [9] were followed. Articles that met the following criteria were included: studies regarding sperm-oviduct interaction and *in-vitro* experimental animal models conducted between 1^st^ January 1990 and 31^st^ July 2024. The population, intervention, comparison, outcome, and study design (PICOS) were followed. (1) population – all *in-vitro* animal experimental studies assessing factors affecting sperm-oviduct interaction (2) intervention – Any non-steroid hormone intervention affecting sperm binding and release (3) comparison – oviduct cells not treated or any other control (4) outcome-effect of intervention, either inhibits sperm binding or enhances sperm binding to the OECs (5) study design – *in-vitro* experimental studies assessing sperm-OECs interaction investigating sperm release and binding to the OECs. The systematic review protocol for the *in-vitro* study could not be registered on PROSPERO, following the website guidelines.

#### Exclusion Criteria

Studies were excluded from the comprehensive literature review if they centered on *in-vitro* studies involving samples from humans, birds, and fish, focused on outcomes other than sperm binding or release from the OECs, steroid hormone supplemented experiments, all *in-vivo* experiments, reviews, letters to the editor, comments, studies not adequately translated into English by google translator, conference proceedings and encyclopedias. Studies published before 1^st^ January 1990 or after 31^st^ July 2024 were also excluded.

### Information sources

The three online databases Web of Science, PubMed, and Google Scholar were used to obtain the articles.

### Search strategy

The inclusion criteria for the search used one phrase from the databases, which read as; (sperm oviduct interactions OR sperm reservoir OR fallopian tube OR oviduct epithelial monolayers OR epithelial explants OR epithelial spheroids AND in vitro techniques OR in vitro experiments AND sperm binding OR sperm release OR sperm density AND in-vitro fertilization AND animal models OR animal studies AND sperm survival OR sperm viability) AND (molecules OR factors) AND (sorted sperm OR sorted spermatozoa OR epididymal sperm). In addition, the bibliography of the selected papers was critically reviewed to identify articles that the automated search could not capture. The online search in the three databases was conducted on 10^th^ August 2024

### Study Selection

The initial review assessed the titles and abstracts of the articles, and any disagreements were resolved through discussion until an agreement was established. Two researchers then independently evaluated the full texts to assess the publications’ appropriateness for inclusion. Any differences between the authors were handled, with the option of discussion.

### Data Extraction

Two authors conducted data extraction independently. Data extracted included the title of the article, objectives, year of publication, authors, semen type used (i.e., fresh, frozen, ejaculated or epididymal, conventional or sexed), species, sample size, factors affecting sperm binding or release from the OEC monolayers, aggregates, or explants. A descriptive synthesis without Meta-analysis was applied because of the inclusion of different methods and protocols in the studies and due to the heterogeneity of the data obtained.

## Results

### Study selection

723 articles were found in the three online database searches and 5 from back referencing. A flow chart of the literature search and study selection is shown in Fig 1. After the removal of duplicates, we reviewed 685 manuscripts. Only 46 articles met the inclusion criteria and were included in the narrative synthesis.

**Fig. 1.**
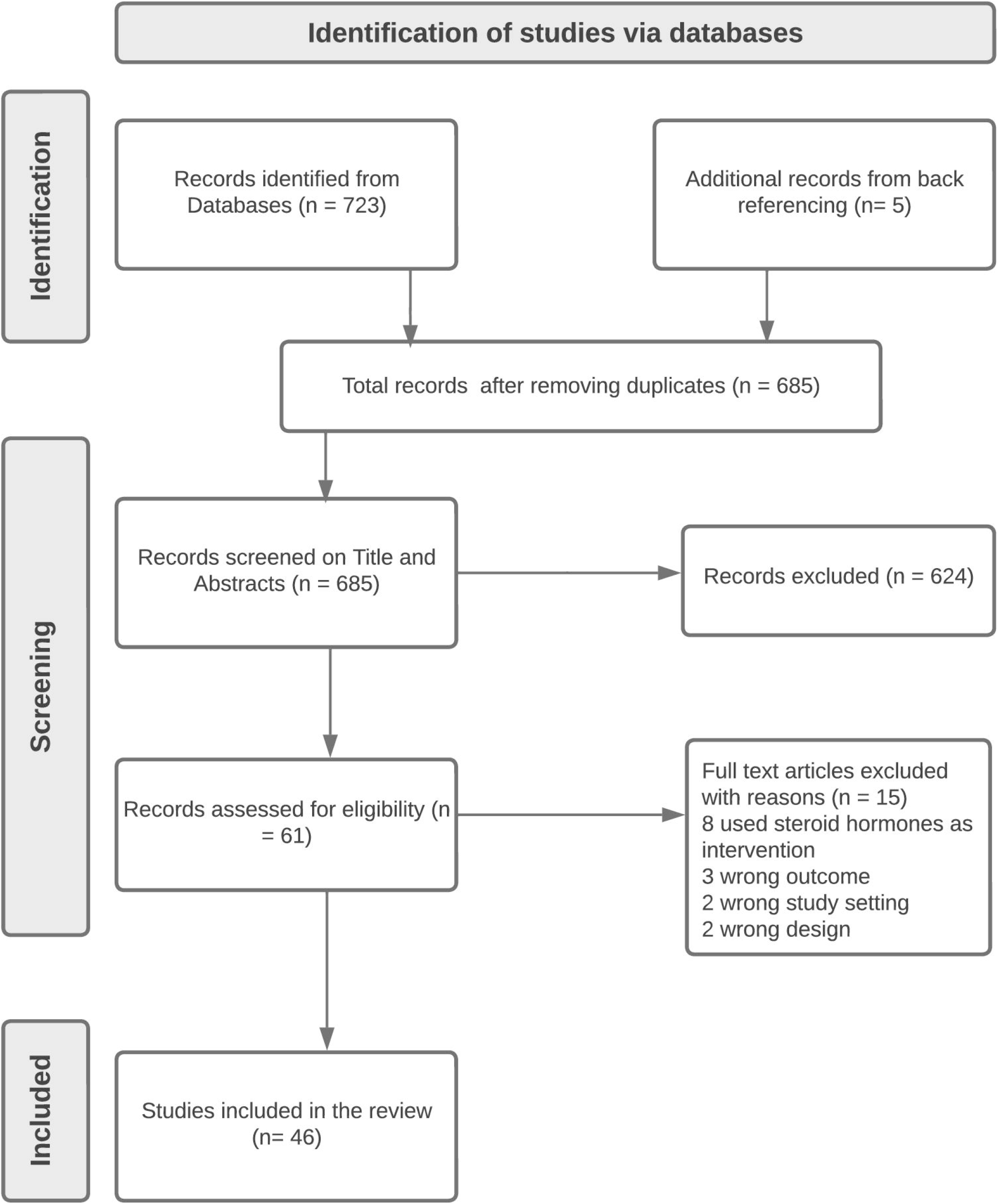
PRISMA flow diagram for the included studies Risk of Bias Assessment.

The risk of bias in the included studies was assessed using the criteria compiled by the National Toxicology Program-Office of Health Assessment and Translation (NTP-OHAT) and adapted for the evaluation of *in-vitro* research [10]. Two independent reviewers evaluated the risk of bias in the included studies. When reviewers could not agree, a consensus was established after discussion. The risk of bias for each included study had six categories assessed and entered in an Excel spreadsheet as definitely high risk, definitely low risk, probably high risk, and probably low risk and later exported to the statistical package for social sciences (SPSS) version 20 for analysis (Fig 2).

**Fig. 2.**
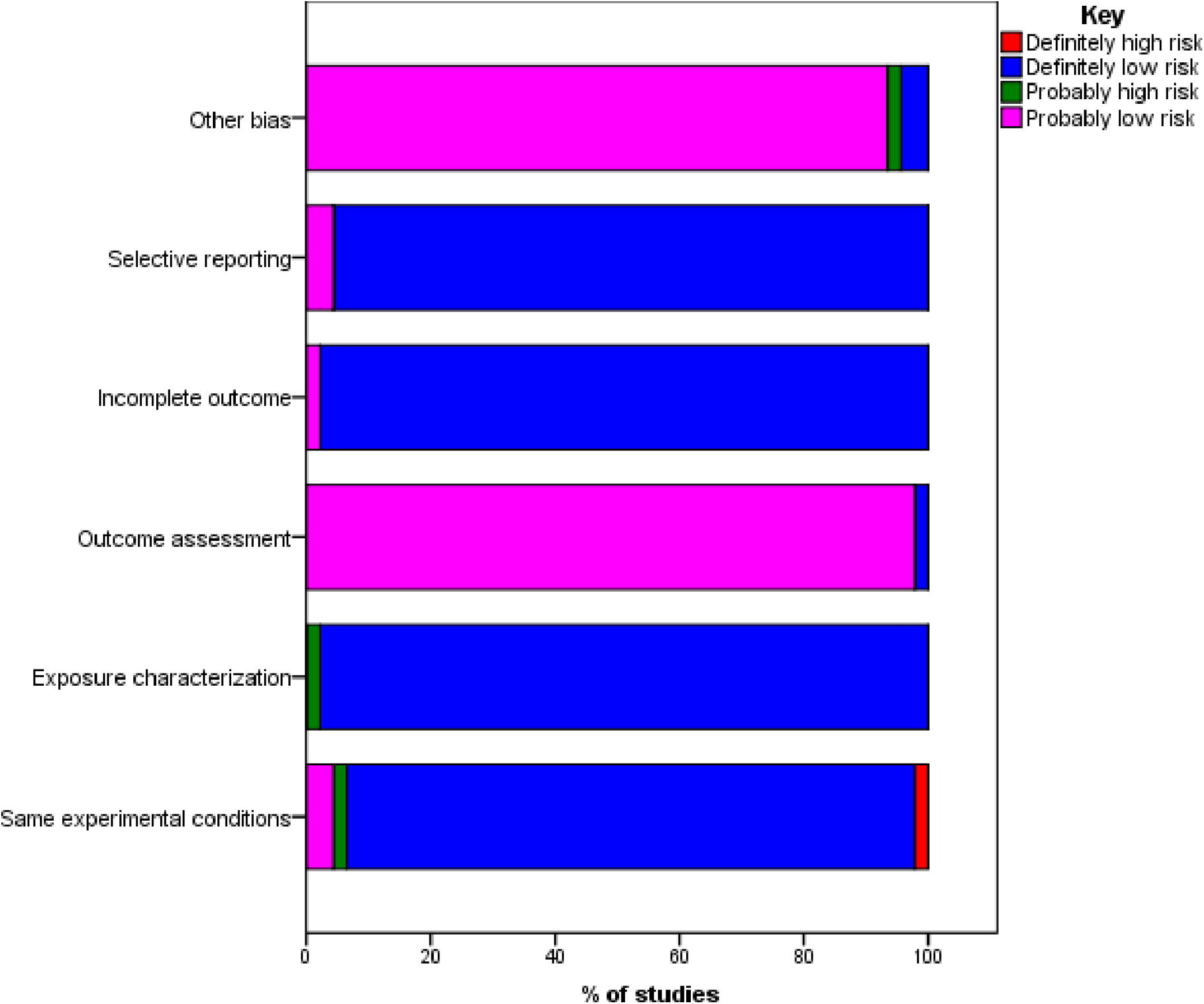
Risk of bias in the included studies using the NTP-OHAT tool.

### Quality Assessment of the Included Studies

The quality of the publications included in the review was evaluated by MCS and MC using a checklist compiled by [11] and blinded to the authors’ names. The assessment revealed that studies did not describe how sample sizes were statistically chosen to improve the power of the studies. Besides, only one study reported blinding of investigators to the allocation of treatment during the sperm binding experiments (Fig 3 and Table 1).

**Table 1:** Quality assessment category and Questions for Fig 3.

| Assessment Category | Question |
| --- | --- |
| I | Is the cell culture passage number reported? |
| II | Is the experimental unit clearly stated? |
| III | Replication of experiment reported? |
| IV | Source of cell lines provided? |
| V | Are cell seeding details reported? |
| VI | Are cell culture characteristics reported? |
| VII | Does the article report how the sample was selected to ensure adequate power? |
| VIII | Random allocation to groups reported? |
| IX | Are investigators blinded to the group allocation? |
| X | Attrition/Exclusions reported? |

**Fig. 3.**
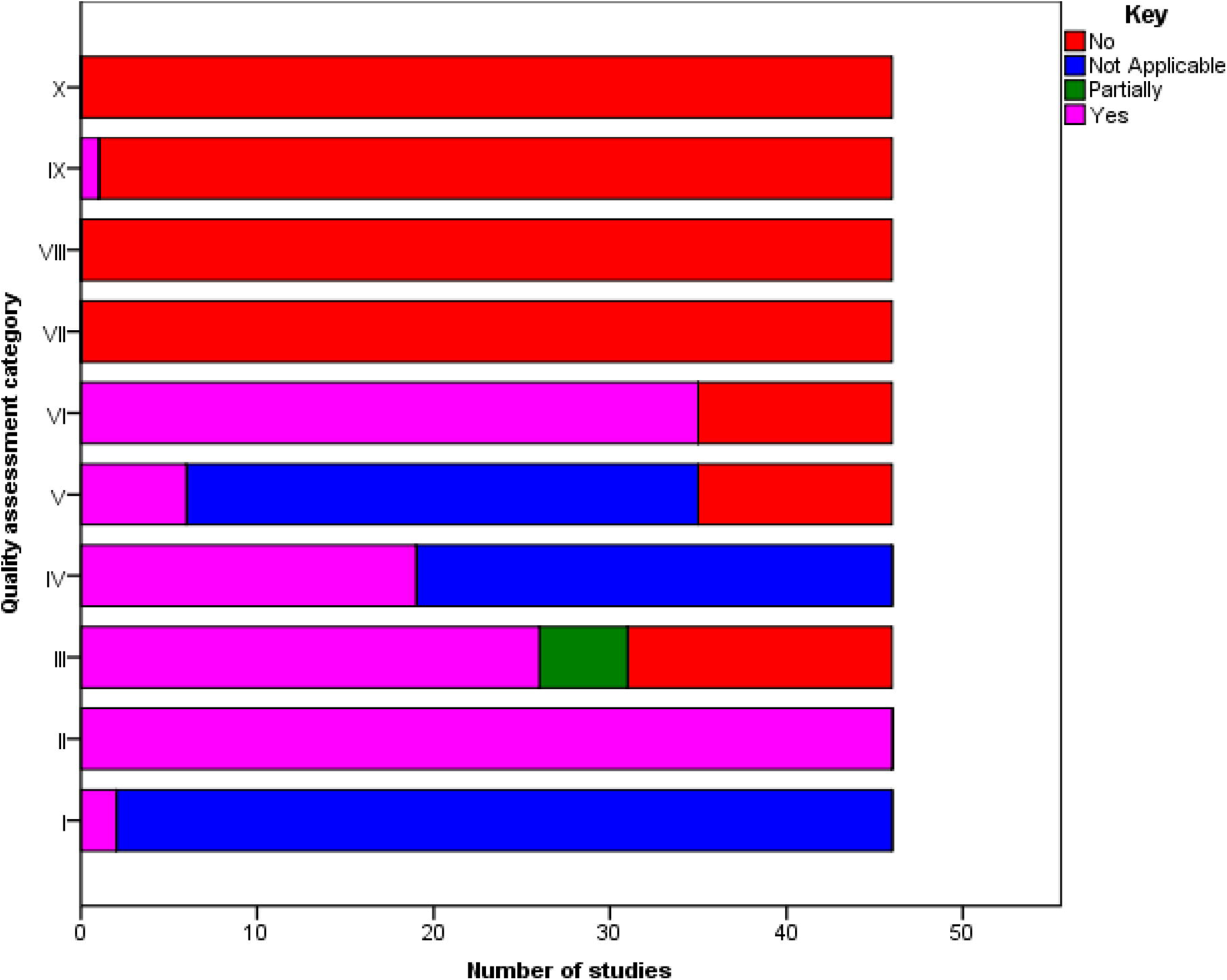
Quality Assessment of the Included Studies.

### Synthesis of Results

#### Biochemical factors affecting sperm release and sperm binding to the oviduct epithelial cells in in-vitro studies

Studies using bovine models (Table 2) showed that sperm binding density to the oviductal epithelium was reduced by the glycosaminoglycan (GAG) heparin [12,13,14,15]. [15] reported no significant difference in the number of sperm bound to the isthmus and ampulla epithelia. This finding suggests that heparin plays a significant role in sperm-OEC interaction, and the subsequent sperm capacitation and release from the oviductal cells that enables sperm to migrate towards the site of fertilization. On the other hand, hyaluronic acid, a non-sulfated GAG increased the binding ability of bull sperm to the epithelial spheroids but there was no significant difference in the sperm binding density to the oviductal spheroids of the preovulatory or postovulatory isthmus and ampulla [12], this signifies the importance of hyaluronic acid contribution to the formation of the sperm reservoir. Another compound, penicillamine enhanced sperm release from the OEC monolayers and OE [13,16]. Notably, penicillamine is not naturally found in the oviductal fluid but is used *in vitro* to improve sperm motility, capacitation, and IVF rates [17,18].

**Table 2:**
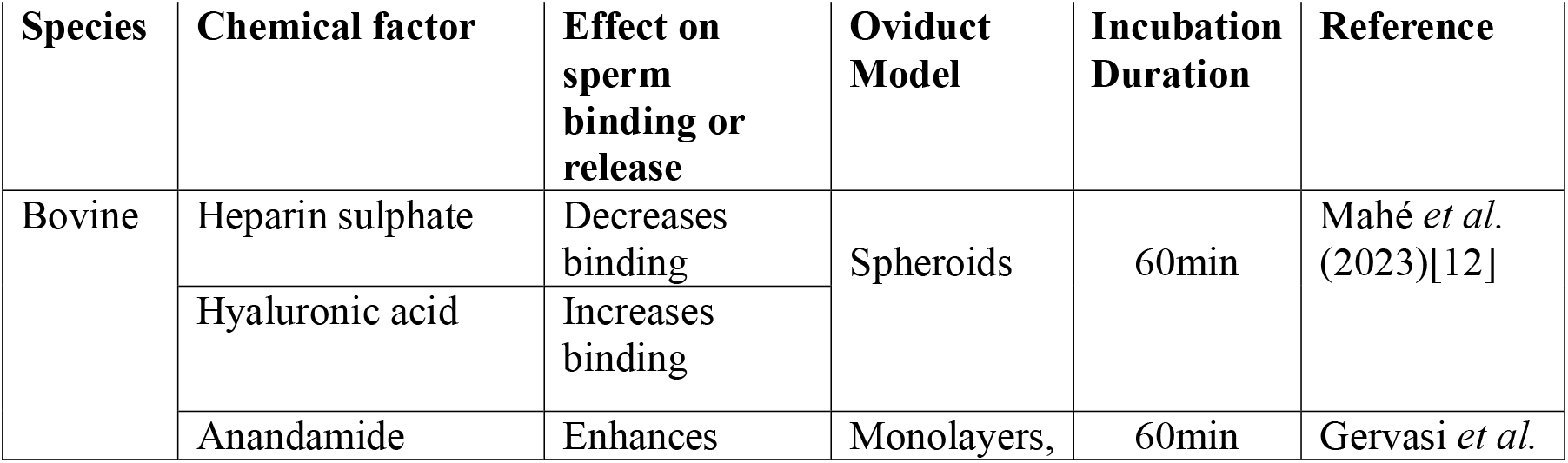
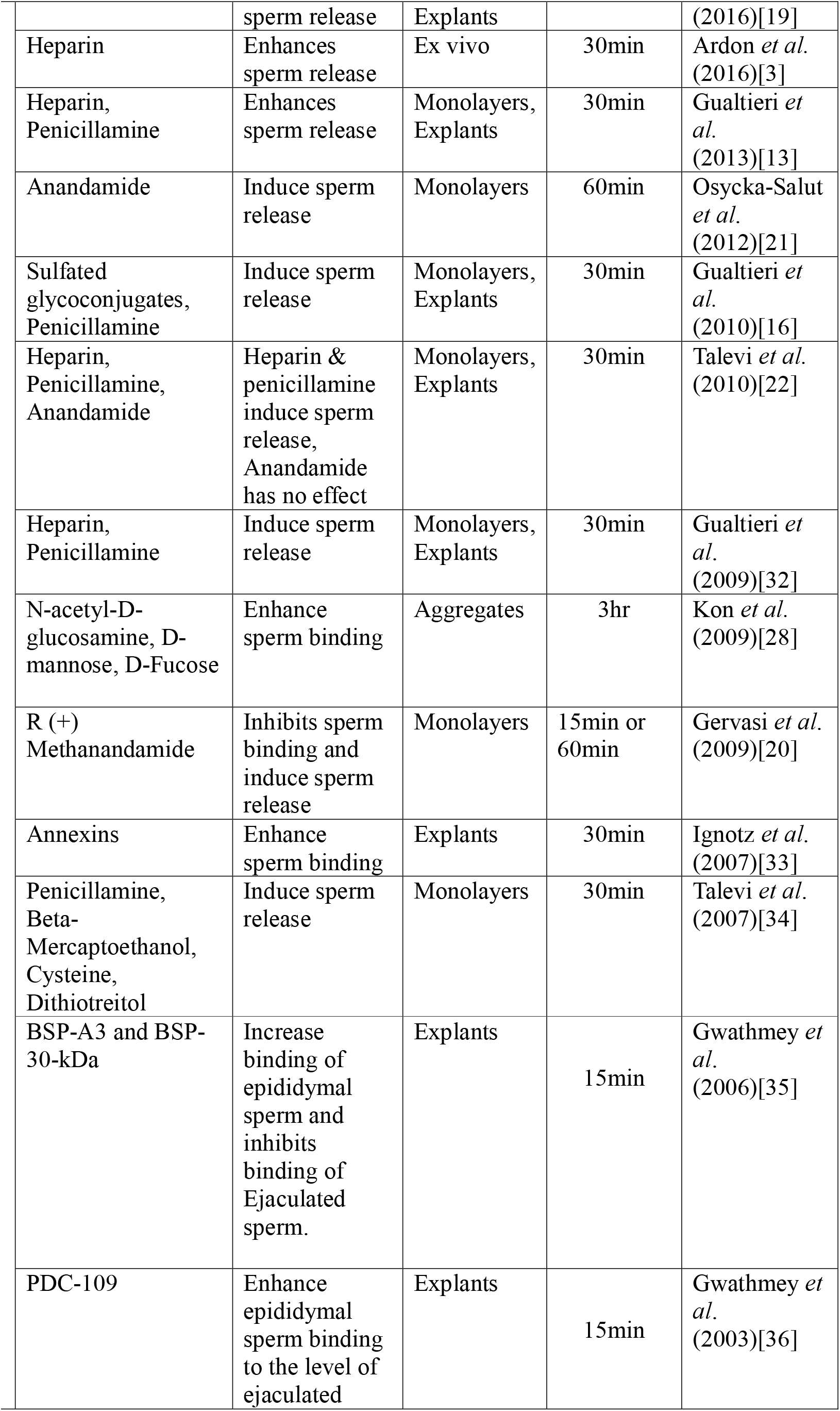
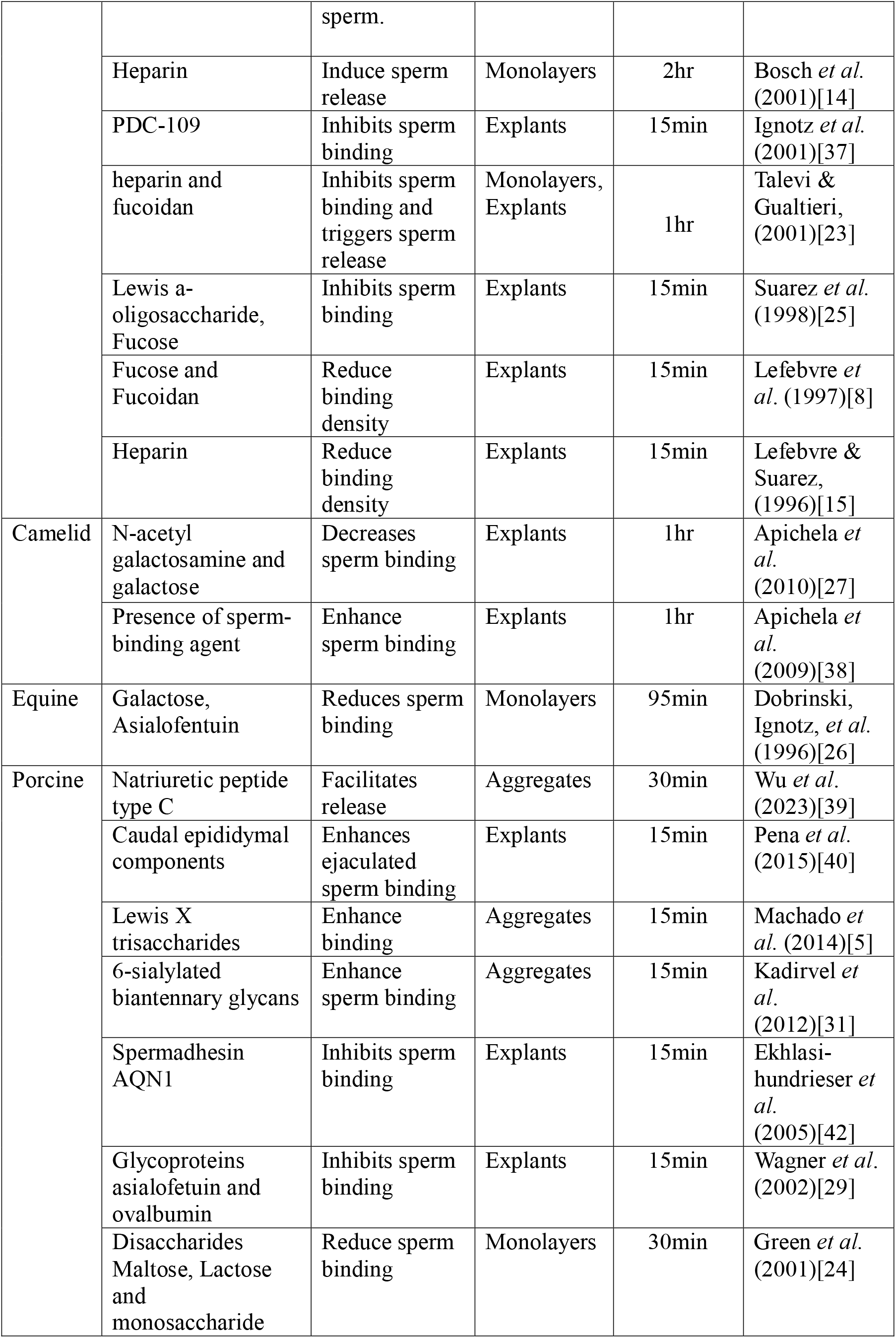
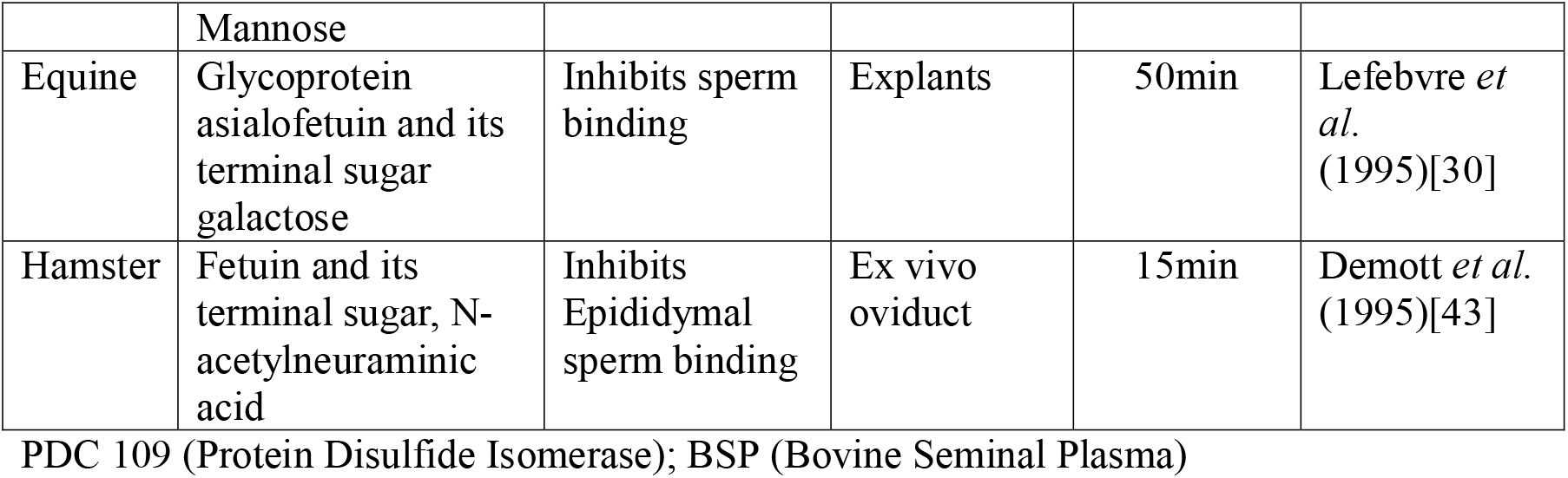
Description of studies assessing the effect of biochemical supplements on sperm binding and release from oviductal epithelial cells *in-vitro* in different animal species.

Another non-steroidal factor inducer of sperm release from the oviduct is the group of molecules belonging to the endocannabinoid. Three studies showed that, anandamide (AEA) facilitated the release of sperm from OEC monolayers and OE through the cannabinoid type 1 (CB1) and the transient receptor potential vanilloid type 1 (TRPV1) activation [19,20] and through the nitric oxide pathway [21]. In contrast, no effect on sperm oviduct binding and release was observed following supplementation with AEA [22]. In these studies, the breeds of bulls and cows for sperm and oviduct collection went unreported respectively. It is possible that variations in experimental conditions, use of sperm from different bulls and differences in expression of the receptors in oviducts obtained from different cows may affect sensitivity, abundance and signalling pathways associated with the receptors.

*In-vitro* models assessing carbohydrates effect on sperm-oviduct interaction reported that; fucose, maltose and mannose, lewis a oligosaccharide fucoidan, and galactose inhibited sperm bull and boar binding to the OEC monolayers and OE [8,23,24,25,26]. The oviductal epithelium’s natural ligands are mimicked by carbohydrates. Consequently, these carbohydrates prevent the sperm from adhering to the epithelial cells by binding to the sperm receptors. likewise, lectin-like interactions are frequently involved in sperm binding to the oviduct. These interactions are likely blocked by excess carbohydrates by saturating the sperm receptors, where sperm surface proteins identify certain carbohydrate structures on the epithelium. Similarly, [27] found that N-acetyl galactosamine and galactose decreased Ilama sperm binding to the OE. Nevertheless, N-acetyl-D glucosamine [28] enhanced bull sperm binding to the OE. On the other hand, asialofentuin inhibited equine and porcine sperm binding to the OEC monolayers and to the OE [26,29,30]. Besides, the terminal galactose residues in asialofentuin mimic the binding sites of sperm on the oviductal epithelium thereby blocking sperm from binding [30].

It is worth noting that monosaccharides are not by themselves on the cell surface but rather components of larger oligosaccharides. The studies that used monosaccharides examined the carbohydrate dependence by attempting to inhibit sperm binding to oviduct cells using very high concentrations of monosaccharides. However, it is worth noting that the physiological relevance of these studies is debatable because inhibition by millimolar concentrations of a monosaccharide may have other effects on cells besides affecting carbohydrate-mediated adhesion. Therefore, future studies must clarify this. Furthermore, [5] and [31] demonstrated that the authentic oviduct glycans bind to porcine sperm and thus participate in the formation of the sperm reservoir.

#### Effect of biochemical supplementation to ejaculated and epididymal semen on sperm-oviduct Interaction

It has been reported that the binding ability of boar spermatozoa develops in the epididymis and that caudal EP components enhance boar sperm binding to the OE [40]. In this study, the binding of EJ spermatozoa to porcine isthmus was higher than that for caudal EP spermatozoa, and EJ boar sperm attached to the isthmus than ampulla epithelium. Interestingly, the seminal plasma protein PDC-109, BSP-A3 and BSP-30-KDa inhibited bull EJ sperm binding to the OE, whereas these proteins enhanced bull EP binding to the level of EJ sperm [35,36,37]. The contrasting effects of PDC-109 on bull EJ sperm and EP sperm binding to the OE could be attributed to differences in the sperm membrane composition and the way PDC-109 interacts with these membranes. Excess PDC-109 might compete with the natural binding sites on EJ sperm, inhibiting their interaction with OE, while it provides EP sperm with the necessary binding capability. Furthermore, the protein annexin was reported to enhance bovine EJ sperm binding ability to the OE [33]. In addition, [16] reported that sulfated glycoconjugates are potent inducers of bovine EP sperm release from the oviductal epithelium. In this study, EP was able to bind to the oviductal epithelium at a lesser extent than frozen-thawed EJ. Likewise, fetuin and its terminal sugar, N-acetylneuramic acid, inhibited hamster EP sperm binding to the oviductal epithelium [43]. This is probably because they may interfere with interactions that are mediated by carbohydrates. N-acetylneuraminic acid and other sialic acid residues found in fetuin can mimic the oviductal epithelium’s natural ligands thus stopping sperm from adhering to the epithelium by competing with these ligands.

### Effect of semen sorting on sperm binding and release from the oviduct

In the studies involving bovine, boar, and ovine sperm-OEC co-cultures (Table 3), it was generally reported that sex-sorting reduces sperm binding to the oviductal epithelium [4,44,45,46]. Therefore, sex sorting and *in-vitro* conditions could significantly affect the expression of adhesion molecules thereby decreasing sperm binding ability to the oviductal epithelium. Interestingly, [47] did not find any differences between the number of ram sperm sex-sorted from frozen-thawed samples and then re-cryopreserved and sex-sorted from fresh ejaculates then cryopreserved binding ability to the oviductal cells. Although not significant the binding ability tended to be lower for sperm sorted from frozen thawed samples than sperm sorted from fresh ejaculates. Therefore, it may be valuable to sort spermatozoa from fresh ejaculates and not from frozen-thawed sperm to prevent further deleterious effects that the cryopreservation and sex sorting processes may cause to sperm.

**Table 3:**
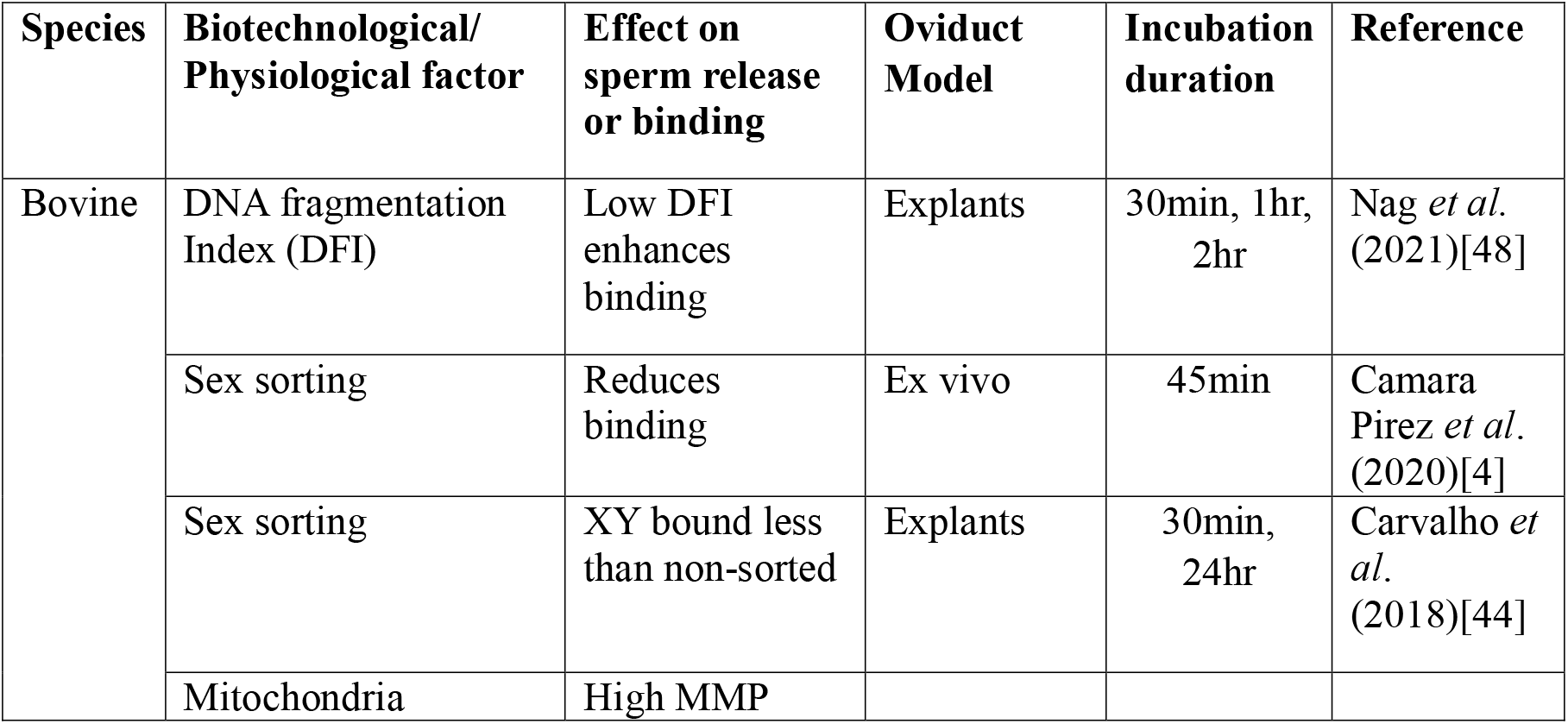
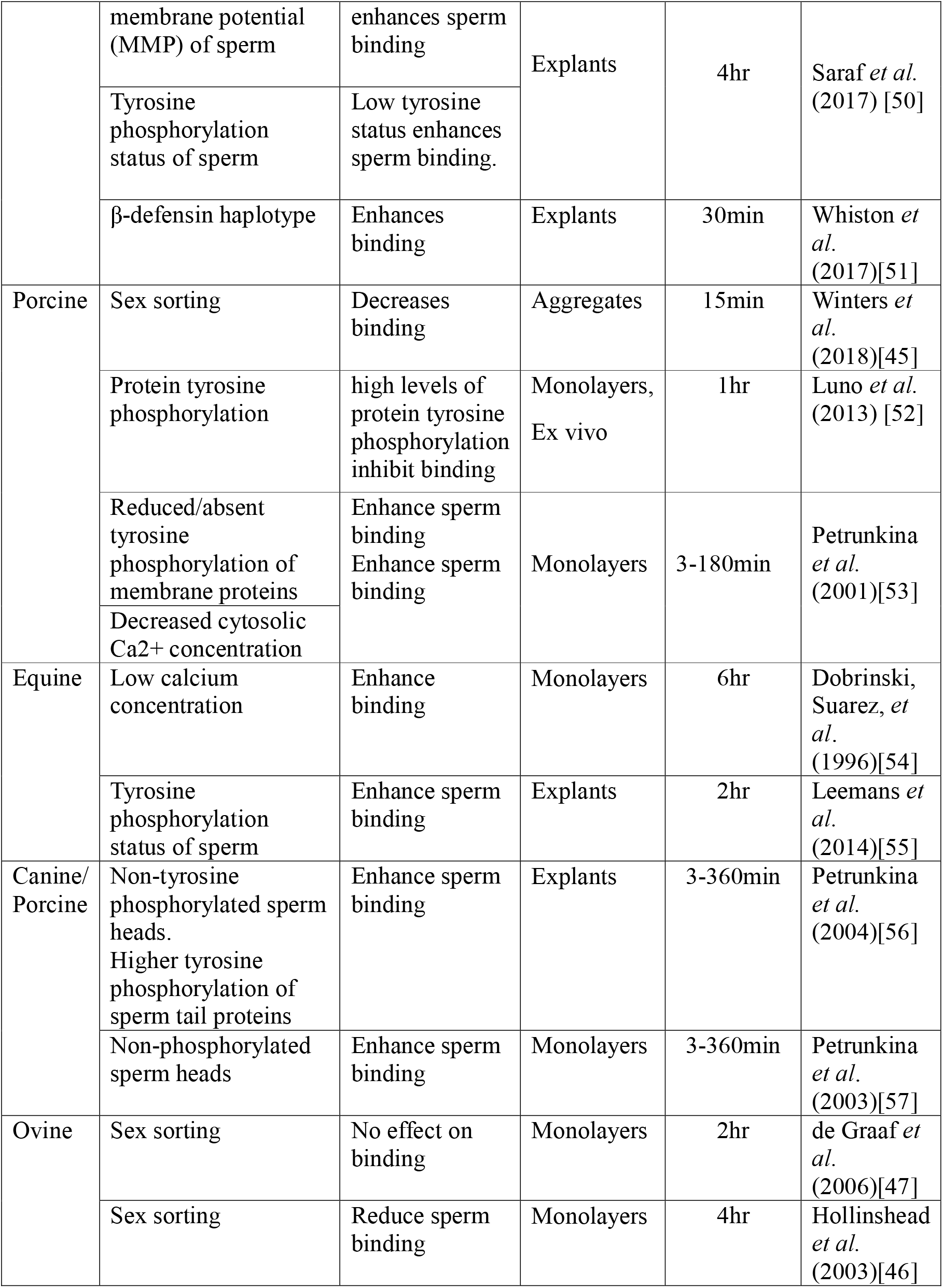
Description of studies assessing the effect of sex-sorting and physiological status on sperm binding and release from oviductal cells *in-vitro* in different animal species.

### Effect of Sperm Physiological Status on Sperm-Oviduct Interaction

It has been reported that bull, boar, equine, and canine spermatozoa with high mitochondrial membrane potential and absent or reduced tyrosine phosphorylation enhanced sperm binding to the OE [50,55,56,57,53]. Likewise, high levels of protein phosphorylation inhibited boar sperm binding to the oviductal epithelium [52].

In addition, bull sperm with low DFI enhanced binding to the OE [48], whereas low calcium concentration of sperm enhanced stallion and boar sperm binding to the OEC monolayers [53,54]. Because the oviduct selects uncapacitated sperm to bind, low tyrosine phosphorylation indicates, low calcium concentration that the sperm are in this pre-capacitated state, making them more likely to bind, while sperm with low DFI tend to have intact plasma membranes, which are crucial for interactions with the oviductal epithelium.

### The oviduct sections used in the *in vitro* experiments

This synthesis shows that various stages of the oviduct sections were used to assess the effect of non-steroid hormone factors on sperm-oviduct interaction in the animal species (Fig 4). The preovulatory stage sections of the oviduct involving bovine, camelid, equine, hamster, and porcine species were used. Further, oviducts sections at different stages of the estrous cycle in bovine and porcine, and in studies involving a heterologous system of canine and porcine sperm-oviduct co-cultures were used. Postovulatory, pubertal, oviducts sections from all diestrus stages and oviducts obtained ipsilateral to ovaries without the corpus luteum used oviducts from bovine, whereas peri-pubertal sections of the oviducts were obtained from porcine. Likewise, pre-pubertal and post-pubertal oviducts from porcine and bovine were used, whereas some bovine studies used non-luteal stage sections of the oviducts. Nonetheless, these studies did not report the influence of the estrous cycle stage of the oviduct sections used on sperm oviduct interaction following supplementation with non-steroid hormone factors. It is essential to evaluate the impact of the estrus cycle stage of the oviducts used in these investigations if our understanding on what happens *in vivo* is to be improved. Moreover, the expression of receptors on the oviductal epithelium is modulated by steroid hormones, including progesterone and estrogen [7,58]. These receptors improve sperm binding and capacitation by interacting with non-steroid compounds such as carbohydrates and glycoproteins. In addition, steroid hormones promote the release of non-steroid substances including glycoproteins and heparin, which foster sperm oviduct interaction and survival [2,59]. Indeed, sperm release from the oviductal reservoir is regulated by a combination of steroid hormones and non-steroid factors, ensuring the optimal timing for fertilization.

**Fig. 4.**
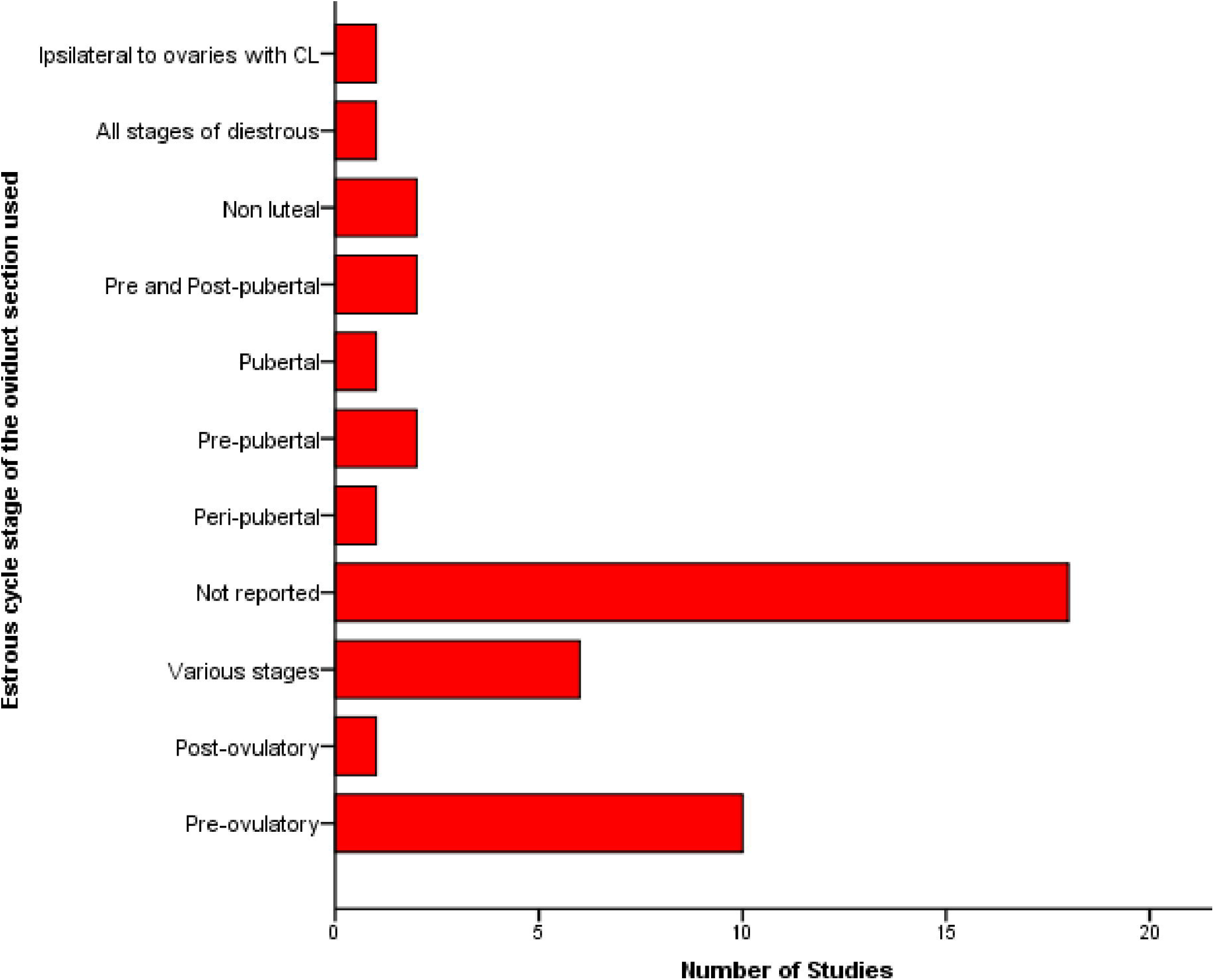
Estrous cycle stage of the oviduct sections used in the *in vitro* experiments.

## Discussion

This review highlights crucial aspects of sperm oviduct interaction, particularly those pertaining to sperm binding and release from the OECs across animal species. The oviduct serves as a critical site for sperm storage and maturation, and the interaction between sperm and oviductal cells is influenced by various biochemical factors, including glycoproteins and carbohydrates. By elucidating these interactions, researchers can improve ARTs, such as TAI and IVF, by optimizing conditions that enhance sperm viability and fertilization success. This understanding may also lead to the development of targeted therapies that improve sperm binding and release mechanisms, ultimately increasing the success rates of reproductive technologies.

Research has established that OEC binding is selective, occurring only with morphologically normal, high-quality sperm that attach via their head region to the oviductal cilia [34]. Our synthesis strengthens the findings that have shown that the carbohydrates and their derivatives enhance sperm binding to the OA [28]. However, contrasting evidence shows that other carbohydrate compounds decrease sperm binding to the oviductal epithelium [24,25,26,27,30]. These contradictory findings likely reflect species-specific differences, highlighting the need for comprehensive cross-species studies to evaluate carbohydrate effects on sperm-oviduct interaction. Such research would enable the development of species-specific protocols to enhance semen sorting and IVF outcomes. During the natural capacitation process that precedes ovulation, changes in sperm surface carbohydrate composition reduce oviductal epithelial adhesion, facilitating sperm migration toward the oocyte [60]. This regulated release mechanism promotes successful *in-vivo* fertilization [60]. Consequently, *in-vitro* sperm-oviduct binding assays incorporating carbohydrate supplementation may serve as an effective tool for selecting high-quality sperm for IVF.

Ejaculated and EP sperm oviduct interaction differences observed could be because, EJ sperm have already undergone modifications during ejaculation, including the incorporation of SP proteins like PDC-109. This protein may saturate or alter the binding sites on EJ sperm, leading to inhibition of binding to OE. On the contrary, while EP sperm typically lack PDC-109, its exogenous addition improves their binding capacity by mimicking the changes that take place after ejaculation. Apparently, the addition of SP proteins PDC-109, BSP-A3, and BSP-30-kDa to EP sperm can improve its fertilizing potential by enhancing various aspects of sperm function [36]. SP proteins can be added to EP in sperm-OEC co-cultures recovered from the caudal epididymis of animals after the slaughter of male animals or the unexpected death of genetically highly valuable animals, and for the preservation of endangered species to improve its fertilizing potential. Strikingly, the conception and farrowing rates were higher in sows and gilts that received insemination using sperm diluted in SP [61]. These factors can contribute to enhanced ARTs by optimizing conditions for sperm selection, capacitation, and fertilization using sperm-OEC co-cultures [62].

Our synthesis showed that sperm bound to the oviduct epithelium had lower levels of tyrosine phosphorylation, implying that the binding process could be a mechanism for selecting sperm with optimal tyrosine phosphorylation levels essential for successful fertilization [52]. Therefore, sperm-OEC co-cultures may be used as alternative methods in the selection of sperm with high fertilizing ability to improve IVF rates. Further, the ability of the sperm to interact with and penetrate the oocyte is strongly associated with the tyrosine phosphorylation of proteins in the sperm head. This alteration improves the sperm’s ability to bind to the zona-pellucida of the oocyte, which promotes optimal fertilization [63].

GAGs such as heparin and fucoidan, and the thiol-reducing agents penicillamine, mercaptoethanol, and dithiotreitol have been shown to play a fundamental role in inducing sperm release from the sperm reservoir. Nonetheless, thiol-reducing agents are not naturally found in the oviduct and thus do not play a fundamental role *in-vivo*, instead they are only used experimentally *in-vitro* to test the importance of protein disulfide bonds in which they induce sperm release from the oviduct epithelium through the reduction of sperm surface protein disulfides to sulfhydryl’s leading to the selection of high-quality and capacitated sperm able to fertilize oocytes [32,34]. The presence of GAGs ensures that only capacitated and motile sperm get released, thus increasing the odds of fertilization [23]. In a study conducted by [64], the supplementation of GAGs to sperm improved the kinematic parameters of sperm, increased the levels of capacitation and acrosome reacted sperm thereby improving the *in-vitro* fertilizing ability of sperm, embryo development rates, and embryo quality and increased the total blastocyst cell number after IVF. Overall, understanding mechanisms that govern sperm release from oviductal cells are critical as they are associated with the selection of high-quality and capacitated sperm with high oocyte fertilizing potential [48]. The oviduct is capable of synthesizing and regulating the concentrations of endocannabinoids along the isthmus and ampulla [22]. Interestingly, fluctuating levels of AEA were found during the estrus cycle in the bovine oviduct [65]. Therefore, understanding the role of the endocannabinoid system in the regulation of mammalian male fertility may provide valuable insights for improving IVF protocols and other ARTs using sperm-OEC co-cultures.

To date, semen sorting has been reported to alter motility, plasma membrane adhesion receptor for the oviduct, membrane destabilization, and capacitation-like changes in mammalian sperm [4,44]. However, the modified sex-sorted sperm behavior, mechanisms, and processes to improve the binding ability of sex-sorted semen to the OEC need further investigation if the conception rates using sex-sorted semen both *in-vitro* and *in-vivo* are to be improved. Our findings showed that sex-sorting reduced the ability of sperm to bind to the oviductal epithelium. Therefore, understanding the sperm structural changes due to the sorting process may help to deduce methods to mitigate these effects leading to increased conception rates following AI *in-vivo* and increased IVF rates. A previous study found the mean blastocyst rates to be low using sexed sperm [66], implying that improvements to the sorting process are required. In our opinion, incorporating specific seminal plasma proteins during the *in-vitro* handling of sex-sorted semen may enhance sperm binding efficiency. Indeed, sex-sorted semen can be enhanced by modifying the sorting process to reduce mechanical stress, thereby improving adhesion molecule expression.

## Conclusion

This review suggests that factors affecting sperm binding and release from the oviduct in different species includes, but are not limited to sex-sorting, glycoproteins, GAGs, carbohydrates, and tyrosine phosphorylation. These factors either reduce or enhance the binding density of sperm to the OECs and may influence the outcomes of ARTs. Moreover, non-steroid factors may provide an alternative to the use of steroid hormones *in-vitro* cultures for the selection of high-quality sperm to be used in IVF program to improve outcomes.

### Limitations

The study has the following limitations. Different methods were used in the studies, which were composed of various components that may affect sperm binding and release differently. Some studies used spherical everted vesicles (epithelial spheroids or explants), while others used monolayers and *ex-vivo* oviducts. This may affect the determination of sperm binding density and the conclusions derived from the studies, making it difficult to compare with the sperm bound to oviductal spheroids. Further, limited publications are assessing sex-sorted semen oviduct interactions, and also wildlife species sperm-oviduct interactions. We suggest that subsequent *in-vitro* experiments should focus on improving the methodological quality of the studies for purposes of enhancing the power of the studies through appropriate sample size determination, random allocation of samples to experimental groups, and ensuring that investigators are blinded to the group allocation during the experiments or when assessing outcomes. Several studies did not report the estrous cycle stage of the oviduct sections used, hence difficult to assess the possible influence of prior exposure of the oviduct to steroid hormones on sperm-oviduct interaction following supplementation with non-steroid hormone factors. Moreover, studies that reported the estrous cycle stage of the oviduct section did not report any effect of the oviduct section on sperm behavior after supplementation with non-steroid factors.

### Future Perspectives

To improve IVF outcomes, further research needs to focus on investigating specific seminal plasma components and their interaction with oviductal glycoconjugates for optimized IVF outcomes. Investigating the effects of different seminal plasma protein concentrations and biochemical supplementation on adhesion molecule expression in sex-sorted versus conventional semen under controlled *in-vitro* conditions is required.

Comparative analyses across species to identify shared versus species-specific mechanisms are required. This can enhance methods for breeding animals and shed light on evolutionary adaptations. Moreover, continued refinement of *in-vitro* models to more closely mimic *in-vivo* conditions, including modulation of the oviduct’s pH and ion concentration, is required. There is further a need to assess and clarify the sperm head protein and molecule composition alteration on the sperm head following the sorting process because these proteins are important in mediating sperm binding to the tubal epithelium. Besides, there is a need to assess and clarify the effectiveness of calcium signaling cascades following the sex-sorted sperm binding to the oviduct epithelium in different animal species. Additionally, there is a need to assess and clarify the impact of the influence of the estrous cycle stage of the oviduct section used to the supplementation with various non-steroid factors on sperm-oviduct behavior in animal species to fully understand *in-vivo* situations. The results derived from such studies may also be used to improve the IVF rates and improve protocols for AI programs.

